# Improved, two-stage protein expression and purification via autoinduction of both autolysis and auto DNA/RNA hydrolysis conferred by phage lysozyme and DNA/RNA endonuclease

**DOI:** 10.1101/2020.01.09.900753

**Authors:** Romel Menacho-Melgar, Eirik A. Moreb, John P. Efromson, Michael D. Lynch

## Abstract

We report improved release of recombinant proteins in *E. coli*, which relies on combined cellular autolysis and DNA/RNA autohydrolysis, conferred by the tightly controlled autoinduction of both phage lysozyme and the non specific DNA/RNA endonuclease from *S. marcescens*. Autoinduction occurs in a two-stage process wherein heterologous protein expression and autolysis enzymes are induced upon entry into stationary phase by phosphate depletion. Cytoplasmic lysozyme and periplasmic endonuclease are kept from inducing lysis until membrane integrity is disrupted. Post cell harvest, the addition of detergent (0.1% Triton-X100) and a single 30 minutes freezer thaw cycle results in > 90% release of protein (GFP). This cellular lysis is accompanied by complete oligonucleotide hydrolysis. The approach has been validated for shake flask cultures, high throughput cultivation in microtiter plates and larger scale stirred-tank bioreactors. This tightly controlled system enables robust growth and resistance to lysis in routine media when cells are propagated and autolysis/hydrolysis genes are only induced upon phosphate depletion.

**Highlights:**

- Autoinduction of both cell lysis and nucleotide hydrolysis
- >90 % lysis and DNA degradation
- Strains are stable to lysis in the absence of phosphate depletion.

## Introduction

*E. coli* is a mainstay for routine expression of recombinant proteins. Recent estimates indicate that over 70% of laboratory studies, reliant on heterologous proteins, utilize *E. coli*. This microbe is commonly used in workflows ranging from high throughput screens, to routine shake flask expression and larger scale fermentations. ^1,2^ In addition, *E. coli* is also used for the manufacturing of proteins at large scale, including the production of over 30% of protein based drugs. ^3,4^ A key challenge to the use of *E. coli* as well as other expression systems where proteins are not secreted, is the recovery of protein from the cell, which routinely requires cell lysis. Common laboratory methods for lysis include: chemical (base or detergents), biochemical (lysozyme) as well as mechanical methods (cell disruptors, french press or sonication), which can not only be tedious and time consuming but yield inconsistent results. ^5–7^ Certain proteins may not tolerate the use of chemical lysis buffers and mechanical methods can lead to incomplete lysis and release of target proteins. In addition, mechanical methods are not amenable to certain workflows such as high throughput screening. ^7–11^ At larger scales, homogenizers are often used to enable more consistent cell lysis, but these units are both costly and add additional steps to commercial processes. ^12,13^ Significant efforts have been made in developing methods for rapid, consistent cell lysis. ^6,8,14–16^, including engineering of *E. coli* strains for autolysis, usually upon induction of one more proteins with lytic activity including: lysozyme, D-amino acid oxidase, muramidase and bacterial phage lysis proteins, which are induced in parallel with proteins of interest and activated after cells are harvested. ^10,11,17–21^

Key remaining challenges with many of these approaches include additional process steps, incomplete lysis or additional induction procedures or vectors. ^10,22–27^ In addition, previous efforts have been focused on cell wall lysis and protein release without consideration of lysate clarification to remove oligonucleotide contamination as well as reduce lysate viscosity. In commercial production after cell lysis, nucleases such as the non specific DNA/RNA endonuclease from *S. marcescens* (Benzonase ®) or alternatives are often used to remove nucleotide contaminants and reduce lysate viscosity to enable easier follow on purification. ^28–30^ the non specific DNA/RNA endonuclease from *S. marcescens* is a small nonspecific extracellular nuclease that is routinely used to hydrolyze contaminating nucleotides during protein purification, and has activity with both double stranded and single stranded DNA as well as RNA. ^31^ An engineered strain of *E. coli* has been reported with periplasmic expression of this endonuclease which auto-hydrolyzes host nucleic acids upon cell lysis, ^32^ but autolysis and autohydrolysis have yet to be combined.

We recently reported strains, plasmid and protocols for the autoinduction of protein expression in stationary phase upon batch phosphate depletion, enabling high protein titers in a very simplified protocol, with no leaky expression. ^33^ To build upon this system, in this work we have further engineered strains with a autolysis and autohydrolysis “module” consisting of lambda phage lysozyme (Lambda R gene) and endonuclease (encoded by the *Serratia marcescens nucA* gene).^28–30^ The expression of a protein of interest as well as expression of the autolysis module are induced upon phosphate depletion coincident with entry into stationary phase, via phosphate regulated promoters (Figure 1a). ^34–39^ These two genes are integrated as an operon into the chromosome in the *ompT* locus (Figure 1b), also deleting this protease, which can lead to improved protein yields. ^40,41^ Tightly controlled (non-leaky) expression and autoinduction media enable a greatly simplified single step process for both high levels of expression and lysate preparation prior to further purification.

**Figure 1:**
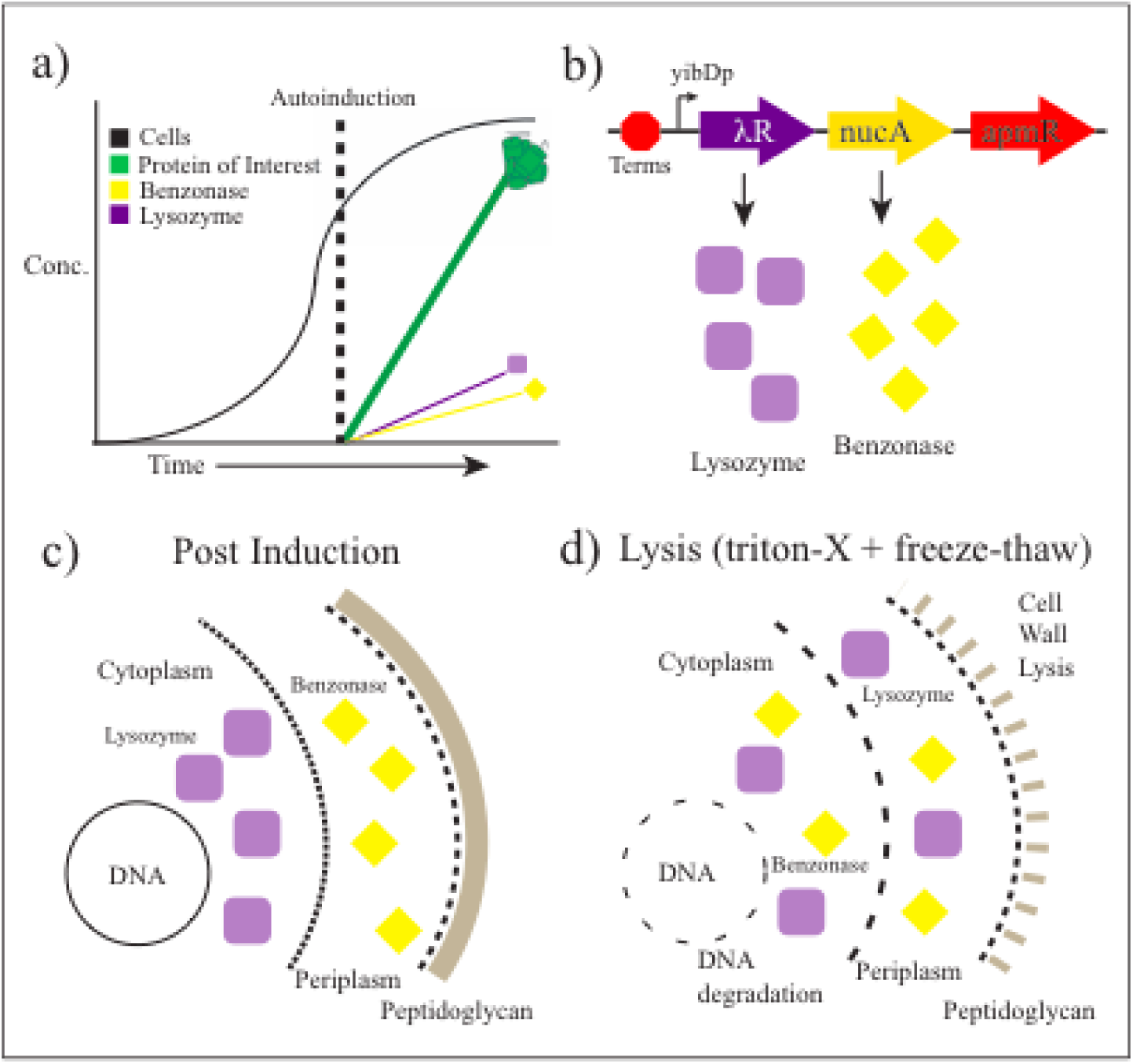
Overview of 2-stage autoinducible autolysis/hydrolysis. a) Cells are grown in optimized media until phosphate is consumed and cells enter a productive stationary phase, simultaneously phosphate regulated promoters are used to induce expression of i) the protein of interest as well as ii) a cytosolic lambda lysozyme and periplasmic endonuclease which are b) integrated as an operon into the chromosome at the *ompT* locus along with an apramycin resistance cassette (apmR) c) both during and post induction lysozyme is separated from the peptidoglycan cell wall by the cell membrane, similarly endonuclease is isolated from cellular DNA. d) Upon the addition of detergents or mechanical stress to disrupt the cellular membrane (in this case triton-X100) the endonuclease and lysozyme can access their respective substrates leading to combined autolysis and autohydrolysis of DNA and RNA.

## Results

### Impact of Autolysis/Hydrolysis Modules on Growth and Protein Expression

After the construction of a modified strain (DLF_R004), with integrated, phosphate regulated lysozyme and endonuclease (Figure 1b), we sought to first evaluate any negative impact these modifications may have on growth and autoinduction of heterologous protein expression. Toward this aim, we evaluated our autolysis/hydrolysis strain as well as its parent lacking any lysozyme or endonuclease for growth and protein expression in autoinduction broth, in the MP2 Labs BioLector ™ (where biomass and protein expression can be monitored). ^33^ Specifically, cells of either strain DLF_R004 (our autolysis/hydrolysis strain) or its parent DLF_R003, were transformed with plasmid pHCKan-yibDp-GFPuv enabling the low phosphate induction of GFP. As can be seen in Figures 2a/b, no significant difference in growth and/or protein expression was observed when the autolysis/hydrolysis module was present. We then further investigated the impact of this module in instrumented bioreactors in minimal autoinduction media, wherein active agitation results in increased shear stresses compared to smaller scale systems. Again, as can be seen in Figure 2c, no significant difference in growth and/or expression was observed indicating strain stability at least to this level of shear.

**Figure 2:**
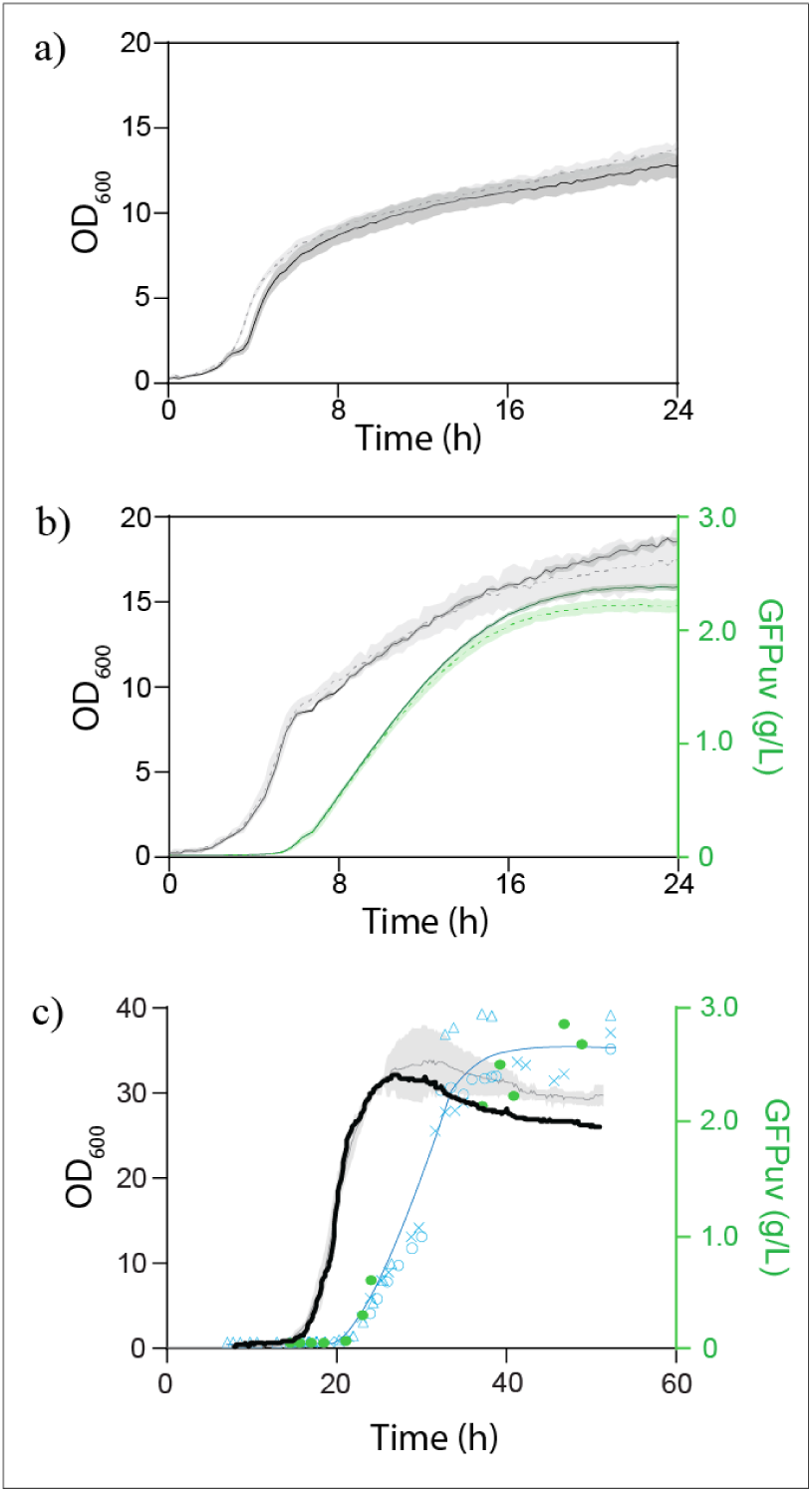
Growth and autoinduction of the autolysis/hydrolysis strain (DLF_R004) compared to a non-autolytic controls (DLF_R002 and DLF_R003). Black and gray line indicate biomass levels (OD 600nm) and blue and green lines and markers indicate GFP levels. Shaded areas indicate the standard error or triplicate evaluations. a) growth of strains DLF_R004 and DLF_R003 in autoinduction broth in the M2P Labs BioLector™. DLF_R003 - dashed line, DLF_R004 - solid line. b) growth and autoinduction of strains DLF_R004 and DLF_R003 both carrying the autoinducible GFP reporter plasmid pHCKan-yibDp-GFPuv, in autoinduction broth in the M2P Labs BioLector™. DLF_R003 - dashed lines, DLF_R004 - solid lines. c) Growth and autoinduction in 1L instrumented bioreactors in minimal mineral salts media. Grey lines and Blue triangles, open circles and squares are three separate control experiments with strain DLF_R002 plus pHCKan-yibDp-GFPuv data taken from Menacho-Melgar et al. ^33^, black line and green circles are results for DLF_R004 plus pHCKan-yibDp-GFPuv.

### Autolysis

After demonstrating equivalent expression with no significant growth defects, we next turned to validate the autolysis behavior of our engineered strain as shown in Figure 3 below. Shake flask cultures were started in autoinduction broth (AB), and the cells were harvested by centrifugation post cell growth and GFP autoinduction. Cell pellets were washed, and triton-X100 was added at 0.1%. GFP release was measured over time by centrifugation and measurement of fluorescence in the supernatant (Figure 3a). The addition of 0.1% triton-X100 was found to be sufficient for the release of ∼ 55% of the total GFP in about an hour. No GFP release was observed either in the control strain or in our autolysis/hydrolysis strain without triton-X100 addition. Increasing triton-X100 levels did not impact protein release (Supplemental Figure S1). To further optimize protein release, we evaluated the impact of a freeze-thaw cycle (Figure 3b) on autolysis. Freeze-thaw is well known to provide cell wall and membrane disruption. As can be seen in Figure 3b, a single 30 minute freeze-thaw after the addition of 0.1% at −20 degrees Celsius, led to >90% release of GFP. Without triton-X100 addition, the same freeze thaw led to only ∼ 10% GFP release (Supplemental Figure S2).

**Figure 3:**
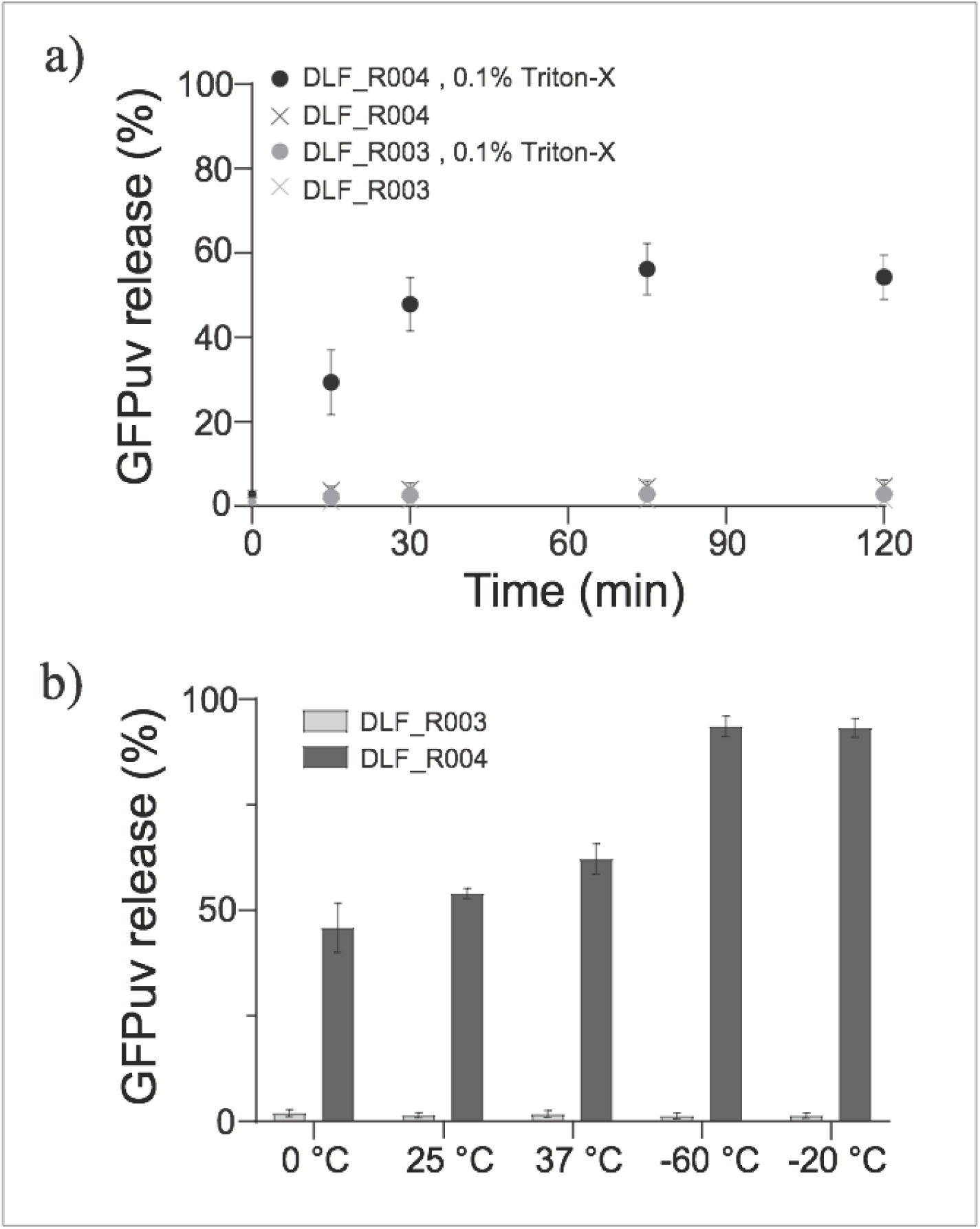
Autolysis and protein release of strains DLF_R004 (autolysis/hydrolysis strain) and DLF_R003 (control). a) Autolysis and GFPuv release as a function of time after the addition of Triton-X100, cells were incubated at room temperature (25 Celsius). b) Autolysis and GFPuv after the addition of 0.1% Triton-X100 and incubation for 30 minutes, on ice (0 Celsius), room temperature (25 Celsius), 37 Celsius, and a 30 minute freeze thaw at either at −60 or −20 Celsius.

### Autohydrolysis

We next turned to validate the autohydrolysis conferred by the endonuclease in our autolysis/hydrolysis strain. To accomplish this we measured DNA/RNA hydrolysis as a function of time during cell autolysis. In the case of hydrolysis, cell lysates were more concentrated to be able to measure differences in DNA concentrations. Cell pellets were resuspended in 1/10th culture volume of 20mM Tris buffer (pH=8.0), plus 2mM MgCl_2_. As a control, EDTA (50mM) was optionally added prior to freeze thawed pellets to inhibit endonuclease. Cell pellets were treated with 0.1 % triton-X100 followed by a single 30 minute freeze thaw. After freeze thaw samples were incubated at 37 °Celsius and samples taken to evaluate hydrolysis. 50mM EDTA was added to samples to inhibit nuclease activity before analysis. Relative levels as well as the size of DNA/RNA were measured both by agarose gel electrophoresis. Results are given in Figure 4 a and b below. DNA hydrolysis, occurs in parallel with autolysis, and visible DNA/RNA was gone within 60 minutes of initiating autolysis. This protocol (with more concentrated lysate) was then evaluated for protein release using GFPuv, results of which are given in Figure 4c, (Refer to Supplemental Figure S3 for DNA analysis) leading to a recommended routine expression and autolysis/hydrolysis protocol for shake flask cultures (outlined in Supplemental Figure S4). Refer to Supplemental Figure S5 for an example lysate generated using this protocol.

**Figure 4:**
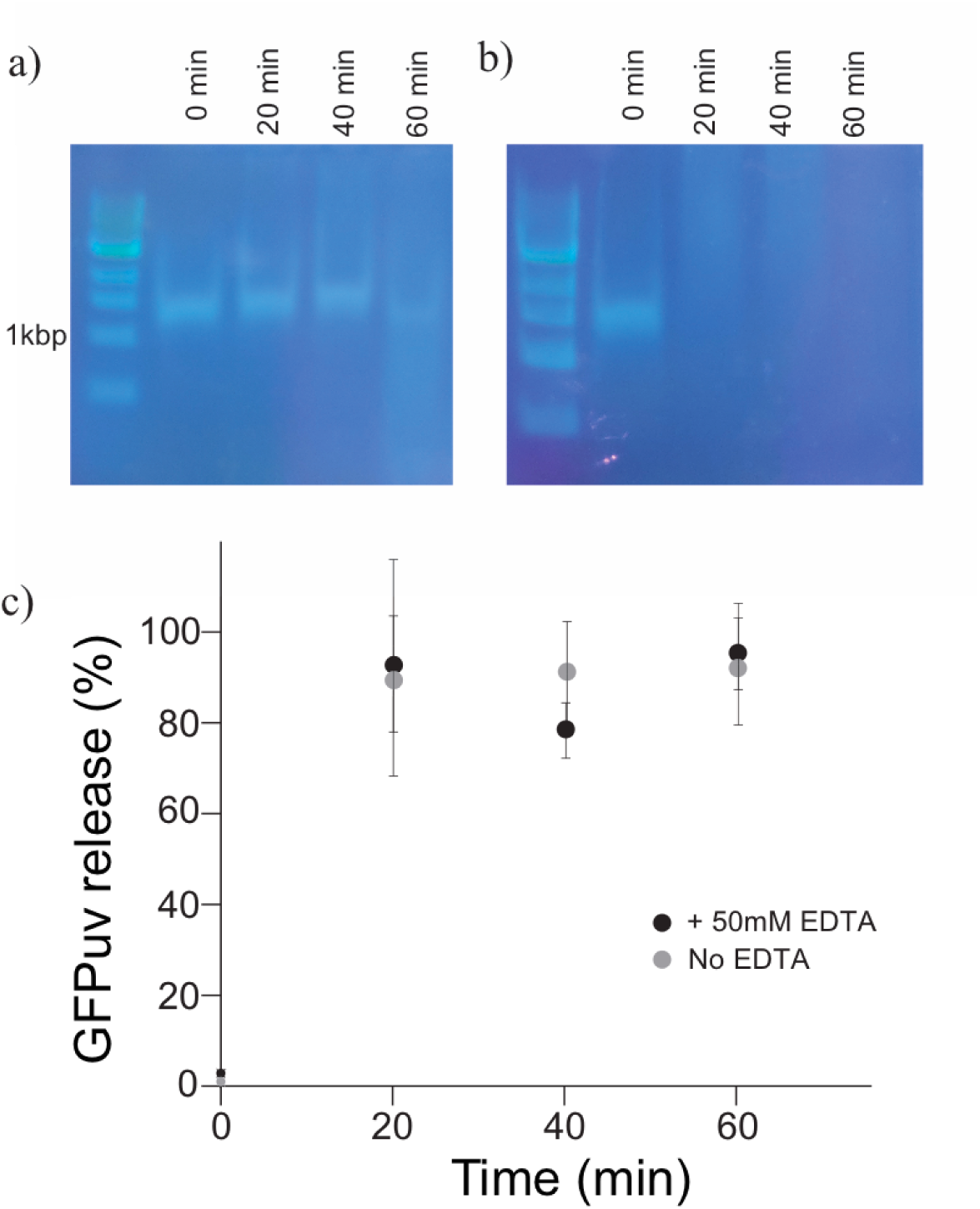
(a,b) Autohydrolysis of DNA/RNA of strain DLF_R004. Time course of DNA/RNA hydrolysis with (a) or without (b) EDTA, which inhibits endonuclease by chelating Mg^+2^. The distinct “band” at ∼ 1500 bp is presumably rRNA and/or sheared genomic DNA. c) Time course of autolysis and GFPuv release in under autohydrolysis conditions using strain DLF_R004 bearing plasmid pHCKan-yibDp-GFPuv,

### High throughput Autolysis/Hydrolysis

To build upon the successful autolysis and autohydrolysis observed in cells harvested from shake flask cultures, we moved to validate this approach with high throughput microtiter based expression. Autolysis/hydrolysis in microtiter plates has the potential to greatly simplify high throughput screening of proteins in crude lysates as well as proteins purified from crude lysates. As illustrated in Figure 5, autolysis and protein release successfully scaled down to MTPs.

**Figure 5:**
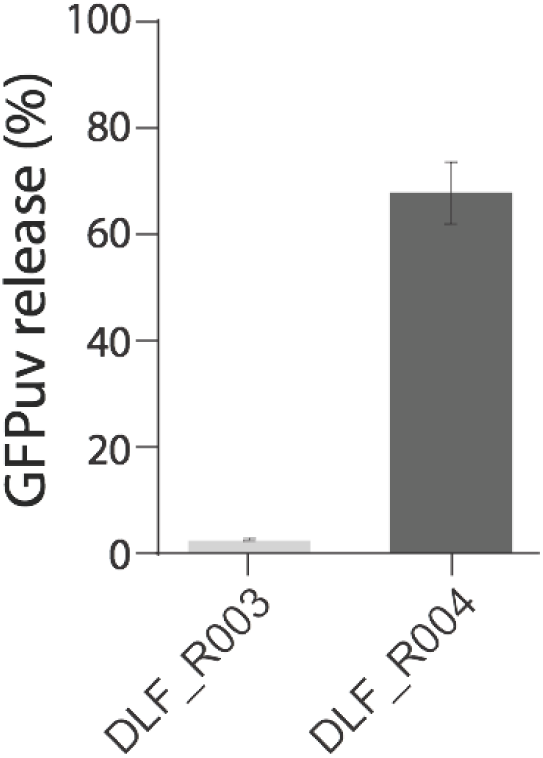
Autolysis and protein release in 96 well microtiter plates.

### Stability of Uninduced Cells

A challenge with several current autolysis strains is sensitivity to free thaw during routine workflows, presumably due to leaky expression of the lysis proteins. And while DLF_R004 has demonstrated stability in autoinduction cultures, we wanted to confirm that autolysis did not occur during routine freeze thaw cycles such as those used in preparing electrocompetent cells where not only are cells frozen and thawed but also thoroughly washed to remove ions including magnesium ions. We tested the stability of electrocompetent cells for both DLF_R004 as well as another well known, readily available autolytic strain of *E. coli*, strain Xjb(DE3) from ZymoResearch. Xjb(DE3) relies on arabinose induction to induce lysozyme and autolytic behavior. ^42^ In addition, the manufacturer recommends that excess magnesium is added to routine cultures to stabilize the cell wall of these cells, which is not feasible when preparing electrocompetent cells. Results of these competent cell studies are given in Figure 6. While strain Xjb(DE3) suffered from unwanted lysis in these studies, DLF_R004, with tight control over expression of lysozyme and endonuclease had increased stability during this process.

**Figure 6:**
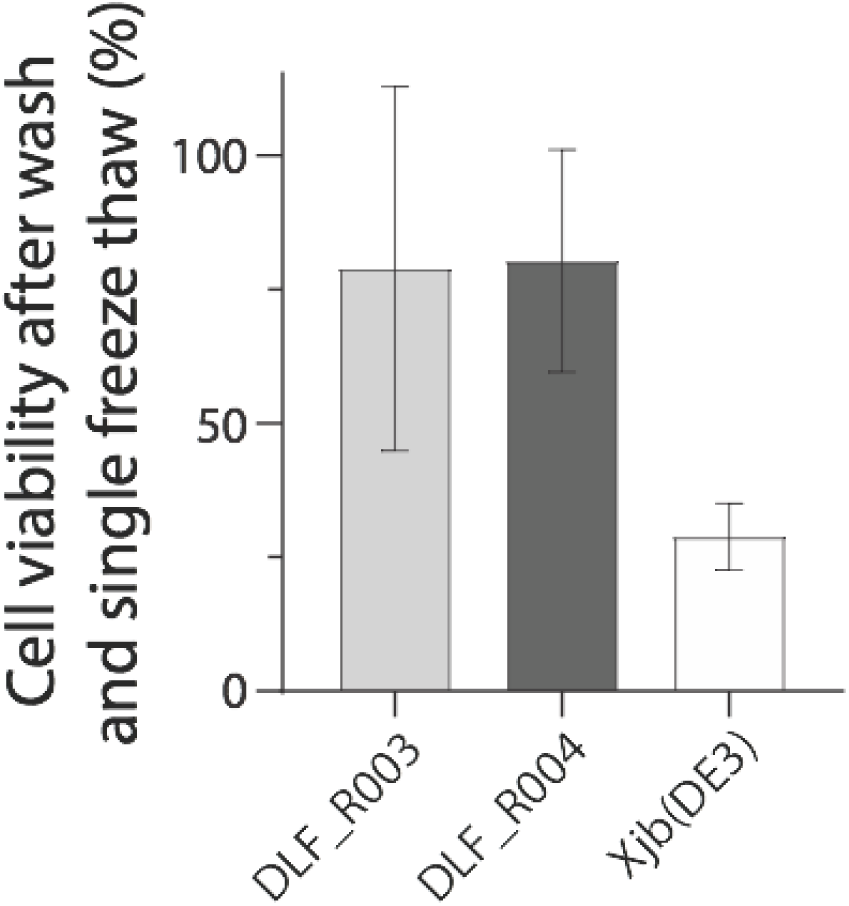
Stability of uninduced strain DLF_R004 (autolysis/hydrolysis strain), DLF_R003 (control) and autolysis strain *E. coli* Xjb. Percent viability was measured after washing with ice-cold water (twice), ice-cold 10% glycerol (once) and a single freeze thaw. Viability was measured as colony forming units after the freeze thaw normalized to colony forming units before freeze thaw, multiplied by 100%.

## Discussion

In this study we have demonstrated the development of an improved strain of *E. coli* for not only autoinduction of protein expression but also of lysozyme and endonuclease enabling combined autolysis and auto DNA/RNA hydrolysis. To our knowledge this is the first combination of these two mechanisms to improve cellular lysis and DNA removal, and an example of the potential benefits of two stage production. ^43–45^ This system enables > 95% lysis and hydrolysis. Due to tightly controlled expression these strains are stable to shear forces in stirred tank bioreactors and even when subjected to freeze thaw cycles in deionized water, with 10% glycerol. Complete autolysis/hydrolysis allows for simplified liquid handling automation, useful in high throughput screening protocols. The mild detergents (0.1% triton-X100) used are also compatible with high throughout SDS-PAGE alternatives including capillary electrophoresis systems, ^46^ and while an optimal buffer was used for optimal autohydrolysis, most buffers are compatible with lysozyme activity and protein release. In commercial production, the autoinduction of endonuclease can remove the need to purchase nucleases for DNA removal and simplify purification and reduce costs.

There are some remaining challenges to the use of the current system in several key applications, due to the use of endonuclease. endonuclease is difficult to inactivate and only denatured under conditions that most likely will impact the activity of any protein of interest. ^47^ As a result, subsequent purification is needed to remove endonuclease. This is likely not an issue for routine shake flask expression or commercial scale production where additional downstream purification steps are expected, but can complicate high throughput screens where lysate assays can be performed and complete purification may not otherwise be needed. This is likely only an issue with high throughput evaluation of enzymes, where retained nuclease activity would be a problem, such as for DNA/RNA modifying enzymes. In these applications additional purification will be needed. Despite these limitations, the method is well suited for routine shake flask expression and protein purification, as well as larger scale production. In addition, the approach may have applicability to the production of other intracellular products beyond proteins including polyhydroxyalkanoates (PHAs). ^48,49^

## Materials & Methods

### Reagents and Media

Unless otherwise stated, all materials and reagents were of the highest grade possible and purchased from Sigma (St. Louis, MO). Luria Broth, lennox formulation with lower salt was used for routine strain and plasmid propagation and construction and is referred to as LB below. Working antibiotic concentrations were as follows: kanamycin (35 µg/mL) and apramycin (100 µg/mL). Autoinduction Broth (AB) and FGM10 media were prepared as previously reported. ^33^

### Strains and Plasmids

Strain Xjb(DE3) was obtained from Zymo Research (Irvine, CA). *E. coli* strains DLF_R002 and DLF_R003 were constructed as previously reported. ^33^ The autolysis/autohydrolysis strain: DLF_R004 was constructed using synthetic DNA. Briefly, linear DNA (gBlock, IDT Coralville, IA) was obtained with the Lamba lysozyme and endonuclease operon driven by a yibDp phosphate controlled promoter, preceded by a strong transcriptional terminator and followed by an apramycin resistance marker (Figure 1b). ^50,51^ The *nucA* reading frame included its native N-terminal secretory signal (‘MRFNNKMLALAALLFAAQAS’). ^28,30^ These sequences were flanked by homology arms targeting the deletion of the *ompT* protease. This cassette (sequence supplied in Supplemental Materials) was directly integrated into the genome of strain DLF_R002 via standard recombineering methodology. ^52^ The recombineering plasmid pSIM5 was a kind gift from Donald Court (NCI, https://redrecombineering.ncifcrf.gov/court-lab.html). *OmpT* deletion and autolysis/autohydrolysis operon integration was confirmed by PCR amplification and sequencing (Genewiz, NC). Plasmid pHCKan-yibDp-GFPuv (Addgene #127078) was constructed as previously reported. ^33^

### Cell Growth & Expression

Shake flask cultures, BioLector™ studies, microfermentations (microtiter plate cultivations) and 1L instrumented fermentations were performed as described in Menacho-Melgar et al.. ^33^ Briefly, batch cultures utilized autoinduction broth (AB Media) and fermentations were performed using FGM10 media. Shake flask expression were performed at 150 rpm in baffled 250 mL Erlenmeyer flasks, with 20 mL of culture.

### Lysis Measurements

DLF_R003 and DLF_R004 strains bearing plasmid pHCKan-yibDp-GFPuv were grown in LB overnight and later used to inoculate 250 mL shake flasks containing AB Media. After 24 hours, cells were harvested by centrifugation at 4000 rpm at 4 °C and resuspended in lysis buffer. Cultures were aliquoted in 1 mL samples. Lysis buffer consisted of either Buffer 1 or Buffer 2. Buffer 1 was used when hydrolysis was not needed and Buffer 2 for autohydrolysis. Buffer 1: phosphate buffer saline pH 7.4 (137 mM NaCl, 2.7 mM KCl, 8 mM Na_2_HPO_4_, and 2 mM KH_2_PO_4_) supplemented with 0.1% Triton-X100 and 1x Halt Protease inhibitors (ThermoFisher Scientific, Waltham, MA). Buffer 2: 20 mM Tris, pH 8.0, 2mM MgCl_2_ supplemented with 0.1% Triton-X100 and 1x Halt Protease inhibitors. Cells were resuspended in 1/10 to ½ the original culture volume (in the case of MTPs). To lyse, cells were incubated in ice (0 °C experiments), preheated heat blocks (25 and 37 °C experiments) or prechilled tube racks (−20 °C and −60 °C experiments) for the indicated time. After lysis, samples were centrifuged at 4 °C at 13 000 rpm for one minute. Fluorescence readings were performed using a Tecan Infinite 200 plate reader in black 96 well plates (Greiner Bio-One, reference 655087) using 200 μl. Samples were excited at 412 nm (OmegaOptical, Part Number 3024970) and emission was read at 530 nm (OmegaOptical, Part Number 3032166) using a gain of 60. Fluorescence values were normalized to complete soluble protein release as obtained from sonicating one sample of each flask using a needle sonicator at 50% power output and 10s/30s on/off cycles for 20 minutes. Under these conditions, we found no more protein release with further sonication.

### DNA Hydrolysis

DLF_R004 and DLF_R004 plus pHCKan-yibDp-GFPuv strains were grown overnight in LB. Overnight cultures were used to inoculate 20 mL of AB, in a 250 mL Erlenmeyer flask at 1% v/v in triplicate. Antibiotics were added as appropriate. Cultures were grown for 24 hours at 37 °C and 150 rpm. Cells were harvested by centrifugation and resuspended in 2 mL of Lysis/Hydrolysis Buffer (20mM Tris, pH 8.0, 2mM MgCl_2_, 0.1% Triton-X100, with or without 50mM EDTA). After resuspension, cells were subjected to a single freeze thaw at −20 °Celsius. Following freeze thaw samples were incubated at 37 degrees Celsius. Samples were taken every 20 minutes, and in the no EDTA reaction, EDTA was added to a final concentration of 50mM. In the case of DLF_R004, samples were clarified by centrifugation and the supernatants analyzed via agarose gel electrophoresis. In the case of DLF_R004 plus pHCKan-yibDp-GFPuv, as GFPuv is also visualized under UV light used to visual agarose gels, after initial lysate clarification and supernatant sampling for GFPuv release, samples were heat denatured at 95 °Celsius for 5 minutes, and then clarified again by centrifugation and the supernatants analyzed via agarose gel electrophoresis.

### Microtiter Plate Expression and Autolysis

96 well plate expression studies again utilized AB media according to Menacho-Melgar et al. using 100 μl of culture volume. ^33^ After 24 hours of growth in AB, cells were harvested using a Vpsin (Agilent) plate centrifuge for 8 minutes at 3000 rpm. Supernatant was removed using a Biotek Plate washer/filler. 50 μl of Lysis Buffer (Buffer 1 above) was added, cells were resuspended by shaking, and placed at −60 °C for 30 minutes. After freezing cells were thawed for 10 minutes at 25 °C, and lysates clarified by centrifugation again using a Vpsin plate centrifuge for 8 minutes at 3000 rpm. 5 μl lysate (supernatant) was collected and diluted 40 fold for analysis of GFPuv levels.

### Strain Stability Measurements

DLF_R003, DLF_R004 and Xjb(DE3) strains were grown overnight in LB. Overnight cultures were used to inoculate 5 mL of LB at 2% v/v in triplicate. The new cultures were grown for 2-3 hours at 37 °C and 150 rpm until 0.6-0.8 OD600 was reached. At this point, samples were taken, diluted 250,000-fold and 50 μL were plated in LB agar plates. The rest of the cells were made electrocompetent by washing twice with 1 mL ice-cold water and once with 1 mL ice-cold glycerol. Cells were then frozen for 2 hours at −60 °C. After thawing, samples were again diluted and plated as described above. Colonies were counted after incubating the agar plates overnight.

## Author contributions

Menacho-Melgar constructed plasmids and strains, performed BioLector™ and shake flask studies. J. Efromson performed fermentations. E. Moreb performed micro-fermentations. R. Menacho-Melgar, J. Efromson, E. Moreb and M. Lynch designed experiments and analyzed results. All authors wrote revised and edited the manuscript.

## Acknowledgements

We would like to acknowledge the following support: DARPA# HR0011-14-C-0075, ONR YIP #12043956, and DOE EERE grant #EE0007563, as well as the North Carolina Biotechnology Center 2018-BIG-6503.

