## Supplementary Materials for "Improved, two-stage protein expression and purification via autoinduction of both autolysis and auto DNA/RNA hydrolysis conferred by phage lysozyme and DNA/RNA endonuclease"

### Supplemental Materials

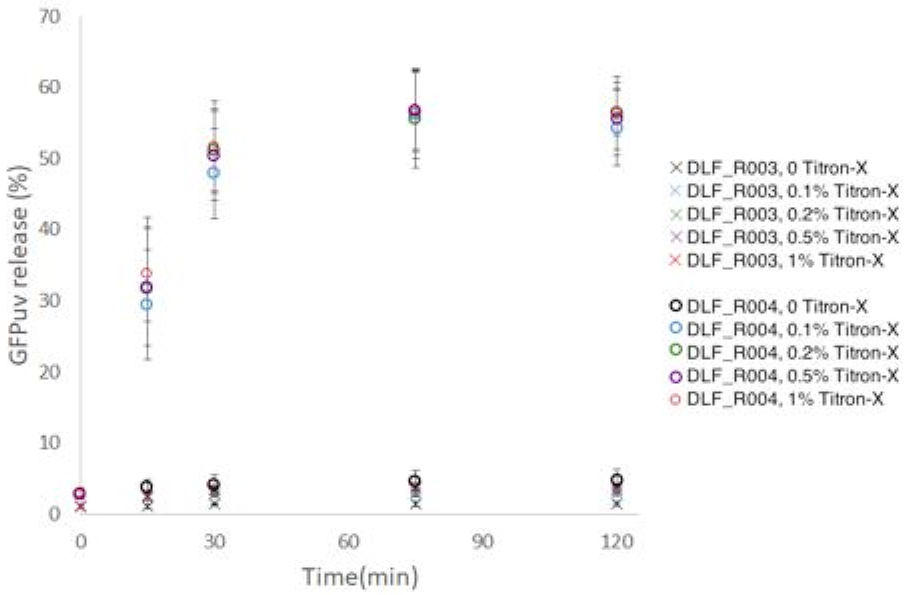


**Figure S1:** Impact of triton level on lysis and protein release. GFP release in strains DLF_R004 and DLF_R003 with the addition of triton X from 0.1% to 1%.

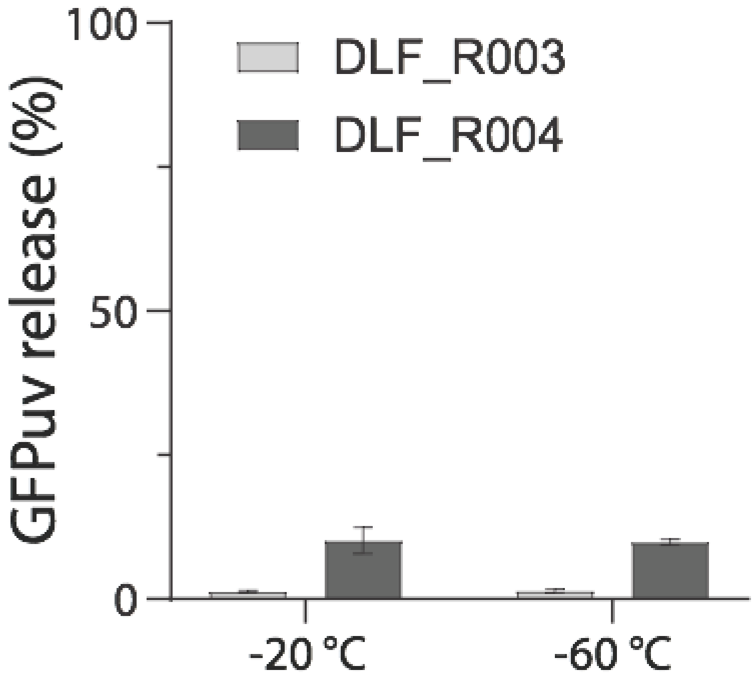


**Figure S2:** Impact of freeze thaw without triton X addition on protein release.

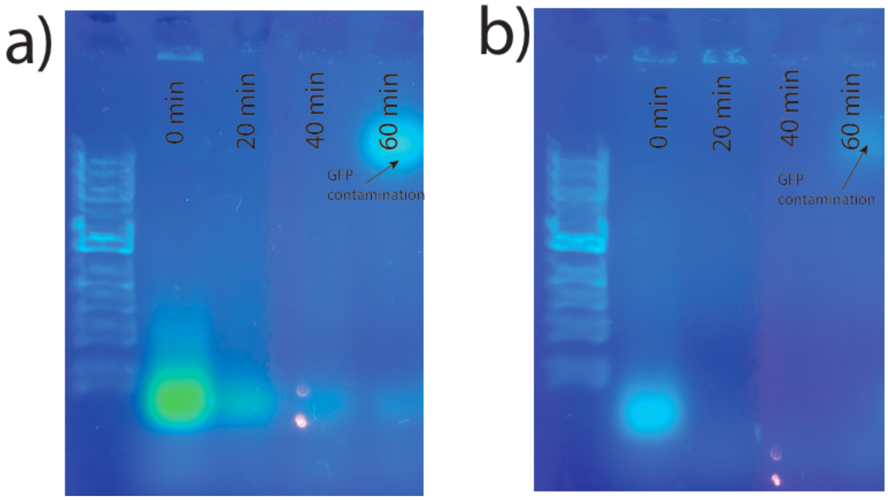


**Figure S3:** DNA hydrolysis in DLF_R004 + pHCKan-GFPuv. a) Agarose gel electrophoresis of heat denatured lysates with EDTA present from the beginning of lysis. a) Agarose gel electrophoresis of heat denatured lysates with active benzonase.

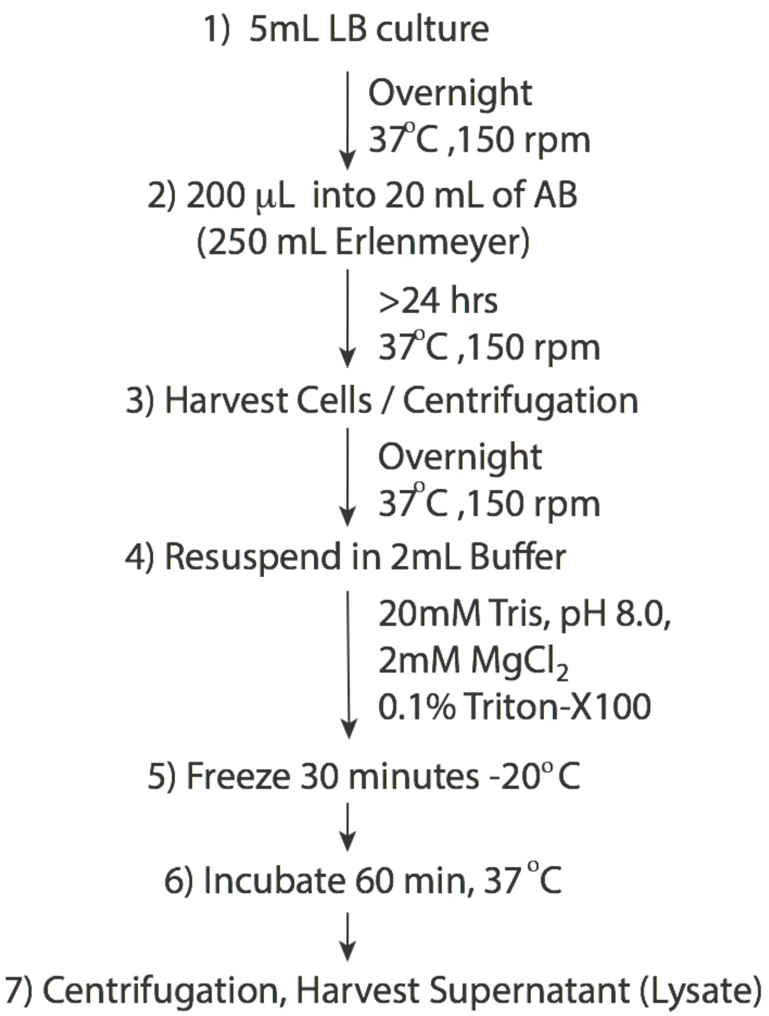


**Figure S4:** Autoinducible Lysis and Hydrolysis Shake Flask Protocol.

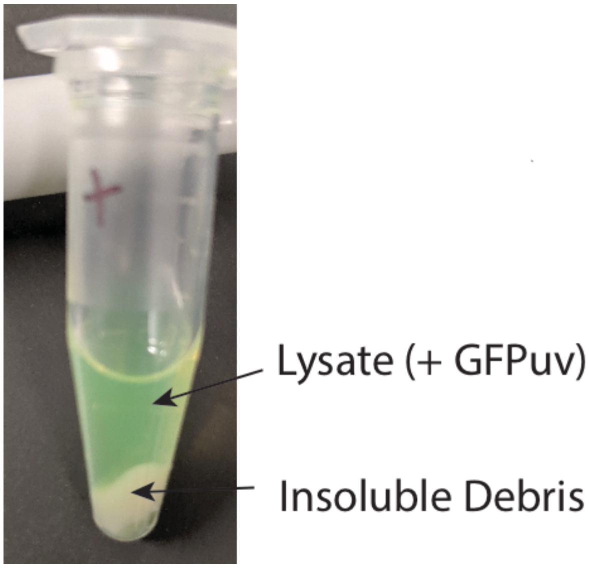


**Figure S5:** Sample of Cleared Lysate.

### DNA And Oligonucleotides used in this study

> delete-ompT-Term-yibDp-Lys-nucA-apmR

AAGAAATTAAAAAACGAAAATACCAGCTATAGCCAGATTGTCACAGAGTGTCGTATGCGTTACGCCGT

ACAGATGTTATTGATGGATAACAAAAATATCACTCAGGTGGCGCAATTATGTGGCTATAGCAGCACGT

CGTACTTTATCTCTGTTTTTAAGGCGTTTTACGGCCTGACACCGTTGAATTATCTCGCCAAACAGCGAC

AAAAAGTGATGTGGtgaAGGGCAAAGCGGAAACGGATAAGACGGGCATAAATGAGGAAGAAATGGCT

CGACCTAGCATAACCCCGCGGGGCCTCTTCGGGGGTCTCGCGGGGTTTTTTGCTGAAAGAAGCTTCAA

ATAAAACGAAAGGCTCAGTCGAAAGACTGGGCCTTTCGTTTTATCTGTTGTTTGTCGCTGCGGCCGGGT

CAGGTATGATTTAAATGGTCAGTAACGGGTCTTGAGGGGTTTTTTGCATATGTGCGTAATTGTGCTGAT

CTCTTATATAGCTGCTCTCATTATCTCTCTACCCTGAAGTGACTCTCTCACCTGTAAAAATAATATCTCA

CAGGCTTAATAGTTTCTTAATACAAAGCCTGTAAAACGTCAGGATAACTTCTGTGTAGGAGGATAATC
TTTTGCTGGAAAAAGGAGATATACCATGGTGGAGATCAATAACCAGCGTAAAGCATTTTTGGATATGT
TAGCGTGGTCTGAAGGGACTGACAATGGCCGTCAGAAAACACGTAACCATGGCTATGACGTGATCGTG
GGCGGTGAATTATTCACAGACTATTCGGACCATCCACGCAAATTAGTAACCCTTAATCCGAAGCTGAA
GTCGACGGGCGCGGGCCGCTATCAACTGCTTAGCCGTTGGTGGGATGCTTACCGCAAGCAGTTAGGGC
TGAAGGACTTTTCGCCGAAGAGCCAGGATGCTGTCGCTCTGCAACAAATTAAAGAGCGTGGGGCTTTA
CCAATGATCGACCGTGGTGACATCCGTCAGGCCATTGATCGCTGCTCTAACATCTGGGCATCTCTTCCA
GGTGCGGGCTACGGGCAGTTCGAGCACAAAGCCGACTCCCTTATTGCTAAGTTCAAAGAAGCGGGAG
GCACGGTTCGCGAGATCGATGTATAAGGATCTAGGAGGGAGATCATATGCGTTTTAATAATAAAATGT
TAGCTCTTGCCGCATTATTGTTTGCAGCCCAAGCGTCAGCGGATACGCTTGAATCTATCGATAACTGTG
CCGTGGGTTGCCCGACCGGGGGTTCGTCTAATGTAAGCATCGTCCGTCACGCCTATACCCTTAACAATA
ATTCTACCACGAAATTCGCGAACTGGGTGGCCTACCATATTACCAAAGATACTCCCGCATCGGGTAAA
ACACGCAACTGGAAGACTGACCCGGCACTGAATCCAGCCGATACATTGGCTCCGGCAGATTACACAG
GTGCAAACGCTGCTCTGAAAGTAGACCGCGGACATCAGGCTCCGCTTGCGAGTCTGGCTGGAGTTAGT
GACTGGGAATCACTGAACTATTTGTCGAATATTACCCCACAGAAATCTGACTTGAATCAAGGCGCGTG
GGCTCGTTTAGAAGACCAAGAGCGTAAACTTATCGACCGTGCAGACATTTCCAGCGTCTATACAGTAA
CAGGTCCTCTGTACGAGCGCGACATGGGAAAGCTTCCTGGTACTCAGAAAGCTCATACAATCCCTTCT
GCATATTGGAAGGTTATCTTTATTAATAATAGCCCGGCTGTCAATCATTATGCCGCCTTCTTATTTGATC
AGAACACTCCCAAGGGGGCTGACTTCTGCCAATTCCGCGTCACTGTAGACGAGATTGAGAAGCGCACG
GGGTTAATCATCTGGGCCGGATTACCTGACGATGTGCAGGCTTCCTTGAAATCCAAGCCGGGAGTTCT
TCCCGAGTTGATGGGTTGCAAAAACTAATAAACCGTGTTGACAATTAATCATCGGCATAGTATATCGG
CATAGTATAATACGACAAGGTGAGGAACTAAACCatgtcatcagcggtggagtgcaatgtcgtgcaatacgaatggcgaaaagccga
gctcatcggtcagcttctcaaccttggggttacccccggcggtgtgctgctggtccacagctccttccgtagcgtccggcccctcgaagatgggccacttggactgatcga
ggccctgcgtgctgcgctgggtccgggagggacgctcgtcatgccctcgtggtcaggtctggacgacgagccgttcgatcctgccacgtcgcccgttacaccggacctt
ggagttgtctctgacacattctggcgcctgccaaatgtaaagcgcagcgcccatccatttgcctttgcggcagcggggccacaggcagagcagatcatctctgatccattg
cccctgccacctcactcgcctgcaagcccggtcgcccgtgtccatgaactcgatgggcaggtacttctcctcggcgtgggacacgatgccaacacgacgctgcatcttg
ccgagttgatggcaaaggttccctatggggtgccgagacactgcaccattcttcaggatggcaagttggtacgcgtcgattatctcgagaatgaccactgctgtgagcgct
ttgccttggcggacaggtggctcaaggagaagagccttcagaaggaaggtccagtcggtcatgcctttgctcggttgatccgctcccgcgacattgtggcgacagccct
gggtcaactgggccgagatccgttgatcttcctgcatccgccagaggcgggatgcgaagaatgcgatgccgctcgccagtcgattggctgagctcaGAACGCC
AACTAAAATTTCCCCGAGGTGAAAATCGCCCCGGGGAATAACTAGCCATTTCAATGTAACAATTAACC
CTTAAAATAAACCCAGAAGGTTATTAACTAAATCACATAGAAAACCATCAATTATAGTATGTATAAAA
TAGGCGACAGCAACCCAATTACAAATTAATGGTTCCAGAATATCACATCAAAAAAAACGCTGTATAAT

ATTATAATTAACATGTAGACAACTTGTAATAAACATTATCAGTCAATTGTTTTGTTTATTCCATCTGTG
ACGCCGATTATTTTCTCAAAATAATGAGATGGCGTGACACCATAATAATCTTTAAATGCACATATGAA
ATATGAAGTACTGTTATAGCCACATTTCTGGGCTACGACATTGATAGAATAAGAGTTTGAAGTTATGA
GTTTTTTTGCATACCTCATCCTAGTATCTCTCAATATTTCAGTAAATGACGTTCCTTCATCCCTTAATCT
TTTTTTTATTAAAC

> OmpT_1000bp_up: GCATTGCTTTTTACCGTATTGTCTAAC

> DLF_R004_seq_conf_2: ATGGTGGAGATCAATAACCAGCGTAAAGC

> DLF_R004_seq_conf_3: TAGCTCTTGCCGCATTATTGTTTGC

> DLF_R004_seq_conf_4: CCTTCTTATTTGATCAGAACACTC

> OmpT_500bp_dn: GATTATTATGGTGTCACGCCATCTC
